# Phenotypic and Genetic Markers of Psychopathology in a Population-Based Sample of Older Adults

**DOI:** 10.1101/601609

**Authors:** Arianna M. Gard, Erin B. Ware, Luke W. Hyde, Lauren Schmitz, Jessica Faul, Colter Mitchell

## Abstract

Although psychiatric phenotypes are hypothesized to organize into a two-factor internalizing – externalizing structure, few studies have evaluated the structure of psychopathology in older adults, nor explored whether genome-wide polygenic scores (PGSs) are associated with psychopathology in a domain-specific manner. We used data from 6,216 individuals of European ancestry from the Health and Retirement Study, a large population-based sample of older adults in the United States. Confirmatory factor analyses were applied to validated measures of psychopathology and PGSs were derived from well-powered GWAS. Genomic SEM was implemented to construct latent PGSs for internalizing, externalizing, and general psychopathology. Phenotypically, the data were best characterized by a single general factor of psychopathology, a factor structure that was replicated across genders and age groups. Although externalizing PGSs (cannabis use, antisocial behavior, alcohol dependence, ADHD) were not associated with any phenotypes, PGSs for MDD, neuroticism, and anxiety disorders were associated with both internalizing and externalizing phenotypes. Moreover, the latent internalizing PGS and the latent one-factor PGS, derived using weights from Genomic SEM, explained 1% more variance in the general factor of psychopathology than any of the individual PGSs. Results support the following conclusions: genetic risk factors for and phenotypic markers of psychiatric disorders are transdiagnostic in European ancestries, GWAS-derived PGSs fail to capture genetic variation associated with disease specificity in European ancestries, and blunt phenotypic measurement in GWAS may preclude our ability to evaluate the structure and specificity of genetic contributions to psychiatric disorders.

## Introduction

Psychiatric disorders impact health, wealth, and wellbeing across the life course (1–3). In the United States, common psychiatric disorders such as Major Depressive Disorder (MDD) are among the top 10 leading causes of disability and injury (4). Among older adults, psychiatric disorders have pronounced effects on physical health and mortality (2,5). Moreover, the 12-month prevalence of having any psychiatric disorder in older adulthood is substantial, with recent estimates of 11.5% (6). As the number of Americans older than 65 years is projected to double in the coming decades (7), more research is needed to understand the presentation and etiology of psychiatric illness in older adults.

### Phenotypic Structure of Psychiatric Disorders

Psychiatric disorders show marked comorbidity across developmental stages (8,9). A robust literature suggests that this comorbidity may be explained by an overarching phenotypic meta-structure that includes separate but correlated internalizing (e.g., depression, anxiety) and externalizing (e.g., substance use, ADHD) factors (10). These comorbidity patterns align with phenotypic differences between internalizing disorders, characterized by elevations in negative affect (11), and externalizing disorders, characterized by behavioral disinhibition (12). Alternatively, a single factor (or a bifactor) model that explains shared variance across all psychiatric disorders has also been supported (10,13), and may emerge in developmental stages where symptoms are less prevalent (e.g., early childhood, older adulthood) (14). Yet examinations of the meta-structure of psychiatric comorbidity have focused primarily on child and younger adult samples. The lack of attention to older adults is a striking omission given the still substantial and impairing rates of psychiatric disorders in this population (2,5,6). Moreover, given clear gender (i.e., greater internalizing symptoms among women, and greater externalizing symptoms among men) and age (i.e., decreasing prevalence of psychiatric disorders across both domains) differences in the prevalence of psychiatric disorders (6,15,16), more research is needed to determine how the meta-structure of psychopathology varies across these demographic groups in older adults.

### Genetic Architecture of Psychiatric Disorders

Genetic risk for psychiatric disorders may also align in a two-factor meta-structure. Twin and family designs suggest that additive genetic risk accounts for the two-factor internalizing-externalizing meta-structure (16,17), and data from genome-wide association studies (GWAS) has been leveraged to identify single nucleotide polymorphisms (SNPs) that are unique to internalizing or externalizing disorders (18). Yet there is also evidence of shared genetic risk across internalizing and externalizing domains (14,19–21), including data from a psychiatric cross-disorder GWAS meta-analysis showing that genetic risk variants are enriched for biological processes core to many psychiatric conditions (21).

As psychiatric disorders are highly polygenic (i.e., resulting from both common variants of small effect, likely to impact many psychiatric disorders, and rare variants of larger effect, possibly unique to certain phenotypes) (22,23), polygenic score (PGS) estimation is one tool that can be used to capture psychiatric polygenicity. A PGS is constructed as sum score of risk alleles that an individual has, weighted by the risk allele effect size from a GWAS in an independent sample (24). Although PGSs are constructed for a specific phenotype (e.g., Major Depressive Disorder), PGS analyses have revealed widespread cross-phenotype correlations (25). For example, a phenome-wide analysis in young adults indicated that a PGS of depressive symptoms was associated with several phobias and generalized anxiety disorder but not externalizing phenotypes, while a PGS for smoking initiation was associated with antisocial behavior but not internalizing phenotypes (26). For researchers studying the etiology of psychiatric illnesses, such widespread associations present a methodological challenge: which PGS best captures genetic risk for a single psychiatric disorder? Given this issue of overlapping genetic risk, quantitative approaches to combining PGSs are needed. Recent methodological innovations have enabled users to construct better-performing PGSs by taking advantage of cross-trait correlations (27– 29). However, such cross-trait “latent” PGSs have only been applied to cohorts aggregated across age-groups (21). In large, population-based samples of older adults, the relative performance of individual PGSs and latent PGSs for specific psychiatric outcomes and general psychopathology is yet unknown.

### Current Study

We assessed the meta-structure of psychopathology in a large population-based sample of 6,216 older adults from the Health and Retirement Study (HRS). We examined whether two-factor phenotypic models fit the data better than one-factor models and further probed the invariance of these models across gender and age. Second, we examined whether there was polygenic specificity in the associations between PGSs for psychiatric (e.g., MDD, ADHD) outcomes and behaviors indexing psychopathology (e.g., cannabis use, antisocial behavior). Next, we implemented Genomic SEM to derive latent PGSs based on the genetic architecture of GWAS summary statistics and present the first analyses of how these latent cross-trait PGSs perform in a large population-based sample of older adults. Based on research in younger samples, we hypothesized that a two-factor model would fit the phenotypic data better than a one-factor model and that PGS-phenotype associations would be hierarchically-organized, such that PGSs for internalizing disorders would be more strongly associated with internalizing outcomes, and PGSs for externalizing disorders would be more strongly associated with externalizing outcomes.

## Methods and Materials

### Sample

Data were drawn from the HRS, a nationally-representative longitudinal panel study of over 43,000 adults over age 50 and their spouses (30). Launched in 1992, the HRS introduces a new cohort of participants every six years and interviews roughly 20,000 participants every two years. Eligible participants for the current study (*N* = 6,003; 58.0% female; Mean age in years [SD] = 67.49 [8.14]) were of genetically European ancestry (i.e., because PGSs were constructed from European Ancestry GWAS) and participated in the Leave-Behind Psychosocial Questionnaire (LBQ) (31) in 2010 or 2012. Participants younger than 51 years were excluded because they were not part of the original sampling frame, as were participants who completed the Leave-Behind Psychosocial Questionnaire in institutional settings, and participants who were born before 1930 (i.e., to address concerns for selective mortality, [30]). Within the analytic sample, 52% earned a high school diploma and 27.4% earned a four-year college degree or higher. Beyond the exclusion criteria listed above, compared to the total HRS, the analytic sample had a higher proportion of women (χ^2^(1) = 11.085, *p* < .001), but did not differ on years of schooling (*t*[38181] = 1.79, *p* > .05). All study procedures were approved by the Institutional Review Board at the University of Michigan.

### Phenotypic Measures

Measures of psychopathology were drawn from the Leave-Behind Psychosocial Questionnaire, a self-reported questionnaire administered to a random 50% of the core HRS participants at each biennial wave during face-to-face interviews (31). A complete wave of data was constructed using the 2010 and 2012 data collections. Depressive symptomatology and drinking frequency were taken from RAND HRS 2010 and 2012 Fat Files (33). All phenotypic data is publicly available through the HRS website (http://hrsonline.isr.umich.edu/).

Measures of internalizing psychopathology included negative affect (34), anxiety symptoms (35), and depressive symptoms (36); externalizing psychopathology was captured by impulsivity (37), trait and state anger (38), and number of drinks per day (Supplemental Table 1).

### Genetic Data and Polygenic Scores (PGSs)

A random subset of the ∼26,000 total participants was selected to participate in enhanced face-to-face interviews and saliva specimen collection (for DNA) in 2006, 2008, 2010, and 2012. Genotyping was conducted by the Center for Inherited Disease Research (CIDR) in 2011, 2012, and 2015. Genotype data on over 15,000 HRS participants was obtained using the llumina HumanOmni2.5 BeadChips (HumanOmni2.5-4v1, HumanOmni2.5-8v1, and HumanOmni2.5-9v1.1), which measures ∼2.4 million SNPs. Individuals with missing call rates >2%, SNPs with call rates <98%, HWE *p*-value < 0.0001, chromosomal anomalies, and first-degree relatives in the HRS were removed. The current paper uses data from unrelated HRS participants of European genetic ancestry (n=9,991) from the genetic data collection years of 2006, 2008, and 2010. Genetic ancestry was determined in a two-stage PCA process wherein the final European American sample included all self-reported non-Hispanic whites that had PC loadings within +/- 1SD of the mean for eigenvectors 1 and 2 in the PC analysis of all unrelated study subjects. PCA was then used again within the European American sample to estimate the top 10 “ancestry-specific” PCs (see Supplemental Methods for more detail).

PGSs of internalizing (neuroticism [37], any anxiety disorder [40], Major Depressive Disorder [38]) and externalizing (alcohol dependence [40], Attention-Deficit Hyperactivity Disorder [41], cannabis use [42], and antisocial behavior [42]) psychopathology were constructed using well-powered, European ancestry GWAS summary statistics (Table 1). If the original GWAS included the HRS, we obtained summary statistics with the HRS sample removed (for more detail, see https://hrs.isr.umich.edu/data-products/genetic-data; [36]). Although GWAS summary statistics are available for other psychiatric disorders (e.g., schizophrenia, bipolar disorder), we did not construct these PGSs because these phenotypes were not measured in the HRS. A PGS for height was included as a negative control (47). To construct PGSs, SNPs in the HRS genetic data were matched to SNPs with reported results in each GWAS (see Supplemental Table 2 for the number of SNPs that contributed to each PGS). As we only used genotyped SNPs (i.e., no imputation) to construct PGSs, we did not trim based on linkage disequilibrium, nor did we impose a GWAS p-value threshold/cut-off for included SNPs (48). The PGSs were calculated as weighted sums of the number of phenotype-associated alleles (zero, one, or two) at each SNP, multiplied by the effect size for that SNP estimated from the GWAS meta-analysis. All SNPs were coded to be associated with increasing disease risk (48). To simplify interpretation, the PGSs were normalized within the European ancestry sample. All analyses in which PGSs were combined with phenotypes included the top 10 ancestry-specific genetic principal components as covariates.

**Table 1.** GWAS summary statistics used to construct polygenic scores

| Construct | GWAS<br>Discovery N | Noverlapping SNPs<br>with HRS<br>genetic data | Citation | Location of GWAS Summary<br>Statistics |
| --- | --- | --- | --- | --- |
| Neuroticism | 170,911 | 1,134,281 | Okbay et al. (2016) | https://www.thessgac.org/data |
| Anxiety | 7,016 cases/<br>14,745<br>controls | 1,079,599 | Otowa et al. (2016) | https://www.med.unc.edu/pgc/results-and-downloads |
| Major<br>Depressive<br>Disorder | 59,851 cases/<br>113,154<br>controls* | 1,340,536 | Wray et al. (2018) | https://www.med.unc.edu/pgc/results-and-downloads |
| Alcohol<br>Dependence | 11,569 cases/<br>34,999<br>controls | 1,170,144 | Walters et al., (2018) | http://www.med.unc.edu/pgc/results-and-downloads |
| Attention-Deficit/<br>Hyperactivity<br>Disorder | 20,183 cases/<br>35,191<br>controls | 1,043,408 | Demontis et al. (2017) | http://www.med.unc.edu/pgc/results-and-downloads |
| Cannabis Use | 162,082 | 1,310,508 | Pasman et al., (2019) | https://www.ru.nl/bsi/research/group-pages/substance-use-addiction-food-saf/vm-saf/genetics/international-cannabis-consortium-icc/ |
| Antisocial<br>Behavior<br>(continuous) | 16,400 | 1,289,915 | Tielbeek et al. (2017) | http://broadabc.ctglab.nl/summary_statistics |

\*Note. The MDD GWAS weights used to construct polygenic scores were based on publicly-available data only and thus, did not include 23andMe data, due to data use agreements. GWAS summary statistics were obtained without the Health and Retirement Study as a contributing sample.

### Genomic SEM

In complement to our analyses using individual PGSs within the HRS, we implemented Genomic SEM to construct latent PGSs (29). Genomic SEM models the genetic covariance structure of GWAS summary statistics and allows for model comparison of different confirmatory factor models (e.g., one factor versus two factor). Single nucleotide polymorphisms (SNPs) can be integrated into the modeling framework to estimate new SNP effects on cross-trait genetic liability, thus allowing for the generation of new PGSs for latent traits. Using the same GWAS summary statistics used to construct PGSs in the HRS, we estimated and compared one-factor and two-factor model of genetic risk for psychopathology. Following Genomic SEM, we construct models of latent PGSs within the HRS, using the same methods described above. It is important to note that traditional confirmatory factor analyses could not be used to evaluate the structure of PGSs because many of the original GWAS included the same participants. Although LD-score regression (49) can be used to determine cryptic relatedness by evaluating the cross-trait LD-score regression intercepts, our analyses revealed substantial sample overlap (Supplemental Figure 1). By contrast, Genomic SEM produces model parameters and test statistics that are unbiased by patterns of shared estimate error across the original GWASs (29).

### Analytic Strategy

All analyses and visualizations were conducted in R Statistical Software (50). To increase generalizability and avoid overfitting the data, the analytic sample (*N* = 6,003) was divided into two random samples of *n* = 3002 and *n*=3001. One dataset (i.e., the “test sample”) was used to estimate phenotypic one-factor and two-factor models using confirmatory factor analyses; the second dataset (i.e., “the hold-out sample”) was used to replicate the best-fitting factor structure. Confirmatory factor analysis is a theory-driven form of structural equation modeling that can be used to capture the shared variance among observed correlated variables to estimate unobserved latent factors (51). The model fitting procedure compares the model implied covariance matrix to the observed covariance matrix, allowing users to compare model fit using several indices. We considered model fit acceptable if the Root Mean Square Error of Approximation (RMSEA) < .06, and the Comparative Fit Index (CFI) and Tucker Lewis Index (TLI) > .90 (52). One-factor and two-factor models were compared using ΔCFI and ΔRMSEA as alternatives to chi-square difference testing, which is sensitive to large sample sizes (53); a ΔCFI > -.01 and ΔRMSEA > .015 indicates significant depreciation of model fit (54,55). All models were estimated using maximum likelihood estimation with robust standard errors in the *lavaan* package (56). Maximum likelihood estimation can be used to account for missing data (in the current study, there was < 5% missing phenotypic data and no missing genetic data), and outperforms other approaches to missing data such as listwise deletion and multiple imputation (57).

The semTools package (58) was used to estimate measurement invariance across gender (1=male, 2=female) and age. To examine invariance across age, we split the sample into three groups: middle age (51 – 64 years), young-old (65 – 74), and old-old (75 – 83). Previous research has documented developmental differences by these age groupings, including environmental effects on depressive symptoms (59), self-rated health (60), and familial social support (61). Increasingly stringent models of invariance across groups are tested: (a) configural invariance – same underlying structure with all parameters freely estimated across groups, (b) metric invariance – invariant loadings across groups, (c) scalar invariance – invariant factor loadings and intercepts across groups, and (d) residual invariance – invariant factor loadings, intercepts, and unique factor variances across groups (62).

Linear regression was used to examine the effects of the individual PGSs and the latent PGSs (estimated using GWAS summary statistics within Genomic SEM) on latent phenotypic factors, controlling for the top 10 ancestry principal components. These analyses were conducted within the hold-out sample only (*n* = 3,003). In large sample sizes, most estimates will be significant at the 95% confidence level. Therefore, we used G*Power (63) to estimate expected effect sizes; assuming 80% statistical power, an alpha error probability of .05, and a sample size of N = 3,003, we are statistically powered to interpret models with an adjusted R^2^ ≥ 0.008.

## Results

Zero-order associations revealed greater within-domain correlations among internalizing phenotypes (.48 < *r* < .64) than externalizing phenotypes (.05 < *r* < .22; Figure 1). However, there were also significant positive cross-domain associations (.16 < *r* < .34). For example, depressive symptoms were positively associated with all the externalizing phenotypes except drinking frequency (Figure 1).

**Figure 1.**
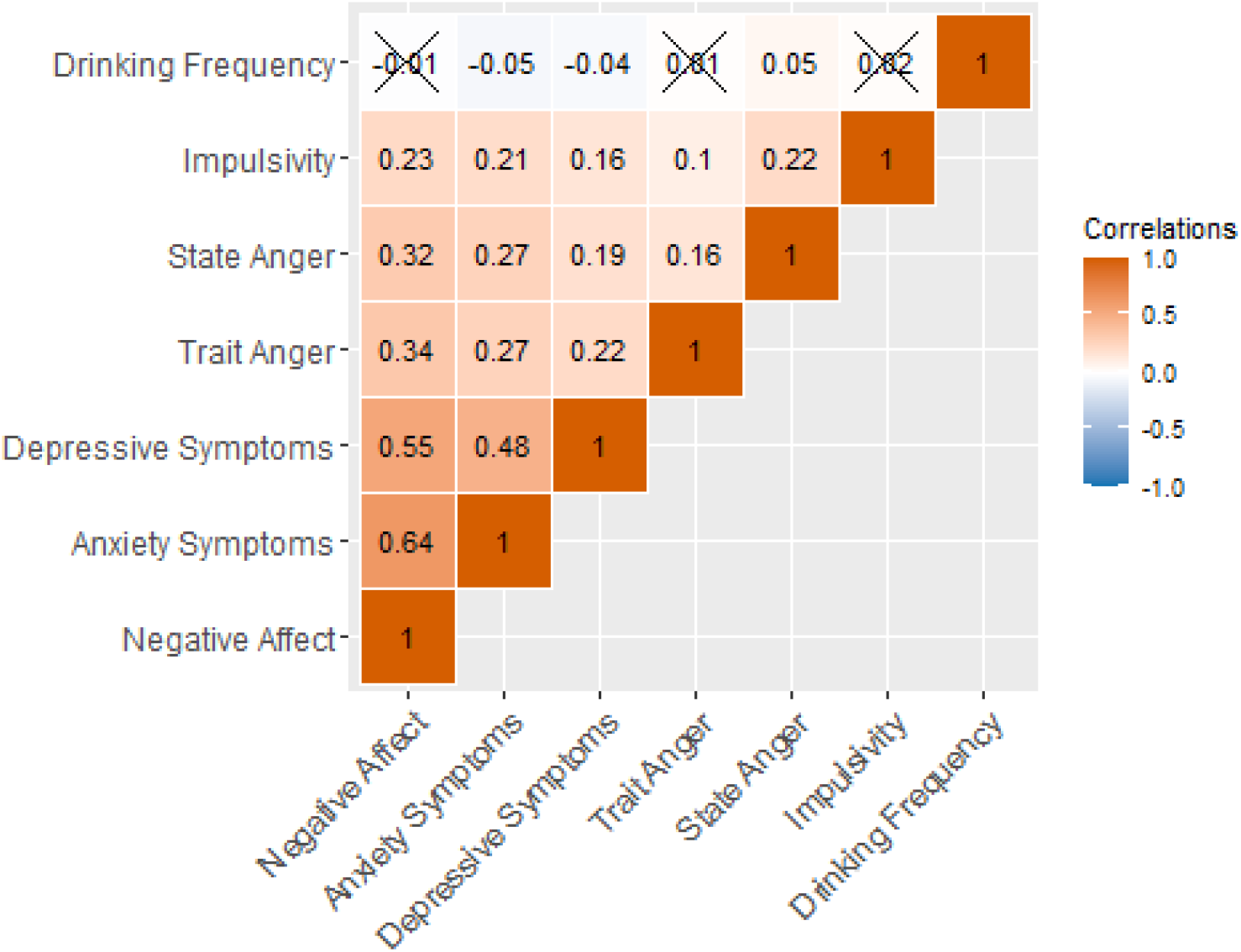
Within- and across-domain correlations among phenotypes in the Health and Retirement Study. *Note*. 5,873 < *N* < 5,965. Associations that were not significant at *p* < .05 are marked with an ‘X’.

### Phenotypic models

Next, we evaluated one-factor and two-factor phenotypic models. In the test sample (*n* = 3,002), drinking frequency loaded negatively on the latent factor(s) and was dropped from subsequent analyses (results available upon request). Figures 2a and 2b display the one-factor and two-factor phenotypic models in the test sample (*n*= 3,002), which both fit the data well. Although the relative fit indices suggested that the two-factor model fit the data better than the one-factor model (i.e., larger CFI and TLI, smaller RMSEA), the ΔCFI and ΔRMSEA were smaller than suggested values (54,55), indicative of equivalent model fit. The association between the internalizing and externalizing latent factors in the two-factor model was very large (*r* = .82), whereas previous work in younger samples reports that the cross-domain correlation hovers around .5 (10). These results suggest that the internalizing and externalizing factors do not represent unique constructs in this sample of older adults in the HRS. Thus, we accepted the one-factor phenotypic model in the test sample. Figure 2c shows the one-factor model in the hold-out sample (*n* = 3,001). The largest loadings for the general factor of psychopathology were negative affect (β = .88, *p* < .001) and trait anger (β = .38, *p* < .001). The general factor explained far more variance in the internalizing indicators (.40 < R^2^ < .78) than the externalizing indicators (.08 < R^2^ < .15).

**Figure 2.**
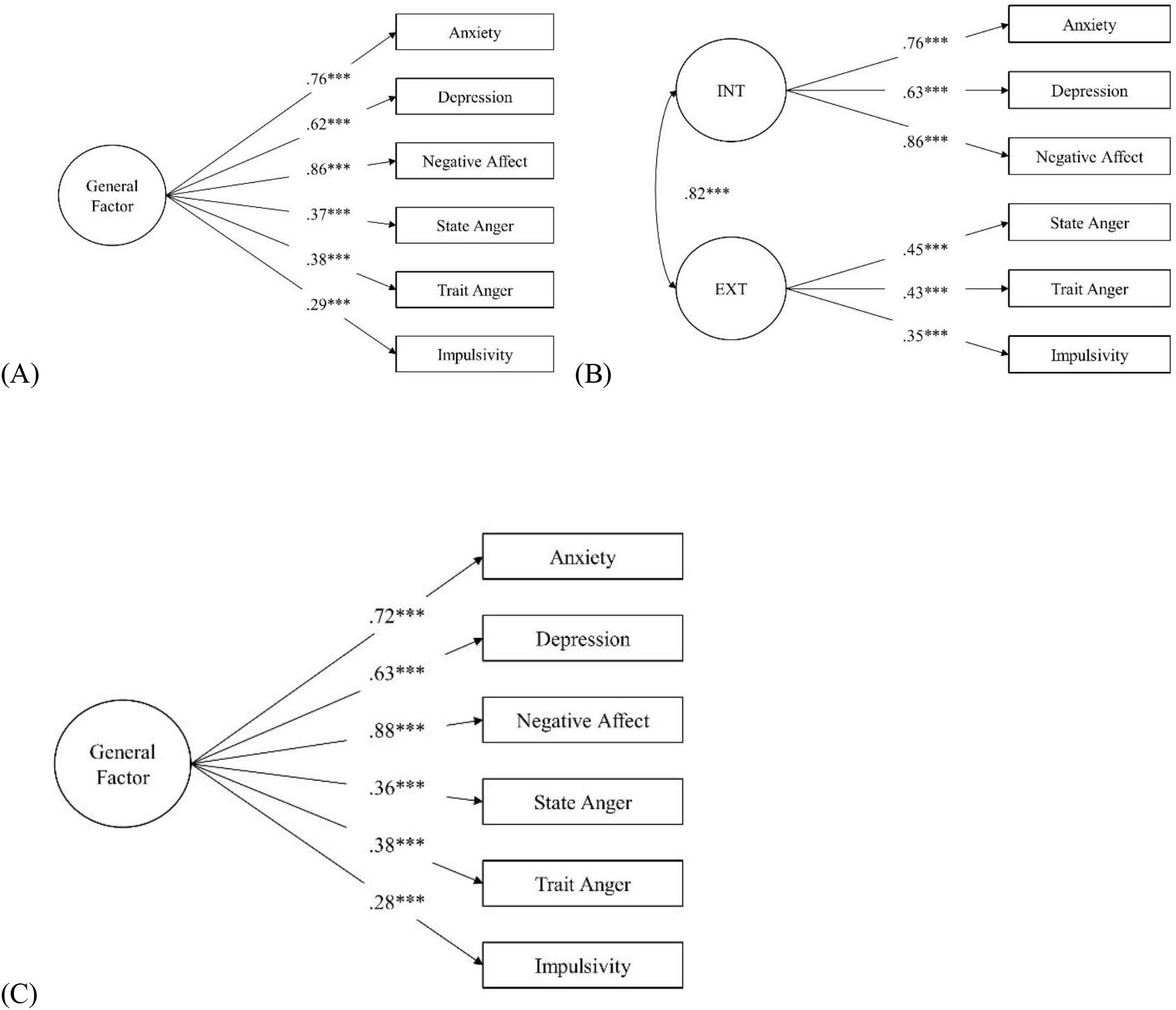
High correlation between internalizing and externalizing factors suggests a one-factor model of psychopathology among older adults in the Health and Retirement Study. *Note*. INT = internalizing; EXT = externalizing. Standardized estimates are shown. (A, B) Confirmatory one-factor (model fit: χ^2^ (9) = 70.37, *p* < .001; CFI = .980; TLI = .967; RMSEA = .052, 90% CI [.041, .064]) and two-factor (model fit: χ^2^ (8) = 46.71, *p* < .001; CFI = .988; TLI = .977; RMSEA = .044, 90% CI [.032, .056]) phenotypic models in the test sample (*n* = 3,002). (C) Confirmatory one-factor phenotypic model in the hold-out sample (*n* = 3,001; model fit: χ^2^ (9) = 61.96, *p* < .001; CFI = .983; TLI = .972; RMSEA = .048, 90% CI [.037, .059]).

We found evidence for metric invariance of the one-factor model of general psychopathology by gender and age group: fixing the indicator loadings to be equivalent across groups did not significantly degrade model fit (see Supplemental Figure 2). As expected, given significant mean-level gender- and age-differences in internalizing and externalizing behaviors (see Supplemental Results), models did not meet criteria for scalar measurement invariance (i.e., equivalent intercepts across groups).

### Polygenic Score Associations with Psychopathology

To address a critical issue in the field, we evaluated polygenic specificity by examining the associations between each PGS and each phenotypic measure, controlling for the first 10 ancestry-specific principle components. Across all phenotypic outcomes, the predictive power of the externalizing PGSs was low (Figure 3a). The only significant association between an externalizing PGS and a phenotypic outcome was a negative association between the PGS for antisocial behavior and impulsivity in older HRS participants. By contrast, the PGSs for neuroticism, MDD, and anxiety were significantly positively associated with anxiety, depressive symptoms, negative affect, and the general latent factor of psychopathology (R^2^ values ∼1%). The PGS for height was not associated with any phenotypic measures.

**Figure 3.**
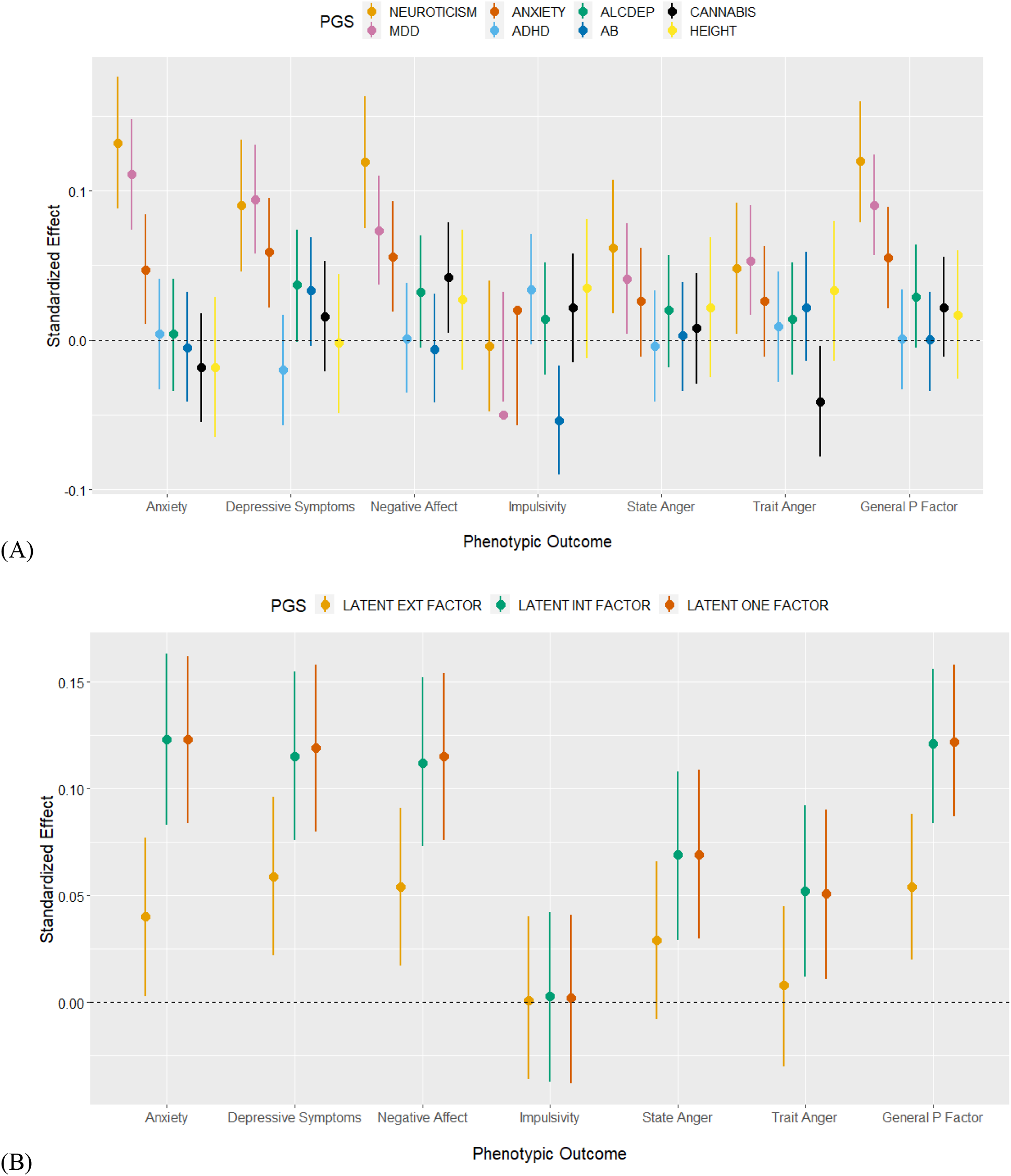
Polygenic scores for internalizing, but not externalizing, disorders are associated with internalizing and externalizing behaviors in the Health and Retirement Study. Note. N = 3,003. Associations between polygenic scores (PGS) and phenotypic outcomes, accounting for the top 10 ancestry principal components. Estimates are unstandardized and error bars are standard errors. (A) Individual PGSs as predictors; (B) Latent PGSs, where SNP weights were estimated using Genomic SEM. In both panels, error bars are standard errors around the estimate.

### Genomic SEM and Latent PGSs for Psychopathology

Genomic SEM was used to fit one-factor and two-factor models of genetic risk for psychopathology, using GWAS summary statistics from well-powered studies of neuroticism [37], any anxiety disorder [40], Major Depressive Disorder [38], alcohol dependence [40], Attention-Deficit Hyperactivity Disorder [41], cannabis use [42], and antisocial behavior [42]. Estimated SNP effects were then used to generate PGSs for latent traits in the HRS sample of older adults. Although both the one-factor and two-factor models fit the data well (Figure 4), model fit comparisons indicated superior model fit of the two-factor model of genetic risk for psychopathology (Δχ^2^ = 30.69(1), p < .001, ΔCFI > .01, lower AIC). Moreover, the cross-trait correlation was *r* = 0.64, indicating that the internalizing and externalizing latent genetic factors, though correlated, capture different underlying constructs. Due to small negative residual variance in the two-factor model, the loading for MDD was fixed to 1. The largest loadings on the latent externalizing factor were alcohol dependence (β = 0.81) and antisocial behavior (β = 0.79). The largest loading on the latent internalizing factor, aside from MDD, was anxiety (β = 0.88). As the model fit for the one factor model was excellent (χ^2^ [14] = 76.762, p < .001, AIC = 104.762, CFI = .962, SRMR = .127), we constructed both the latent one factor PGS and latent internalizing and externalizing PGSs.

**Figure 4.**
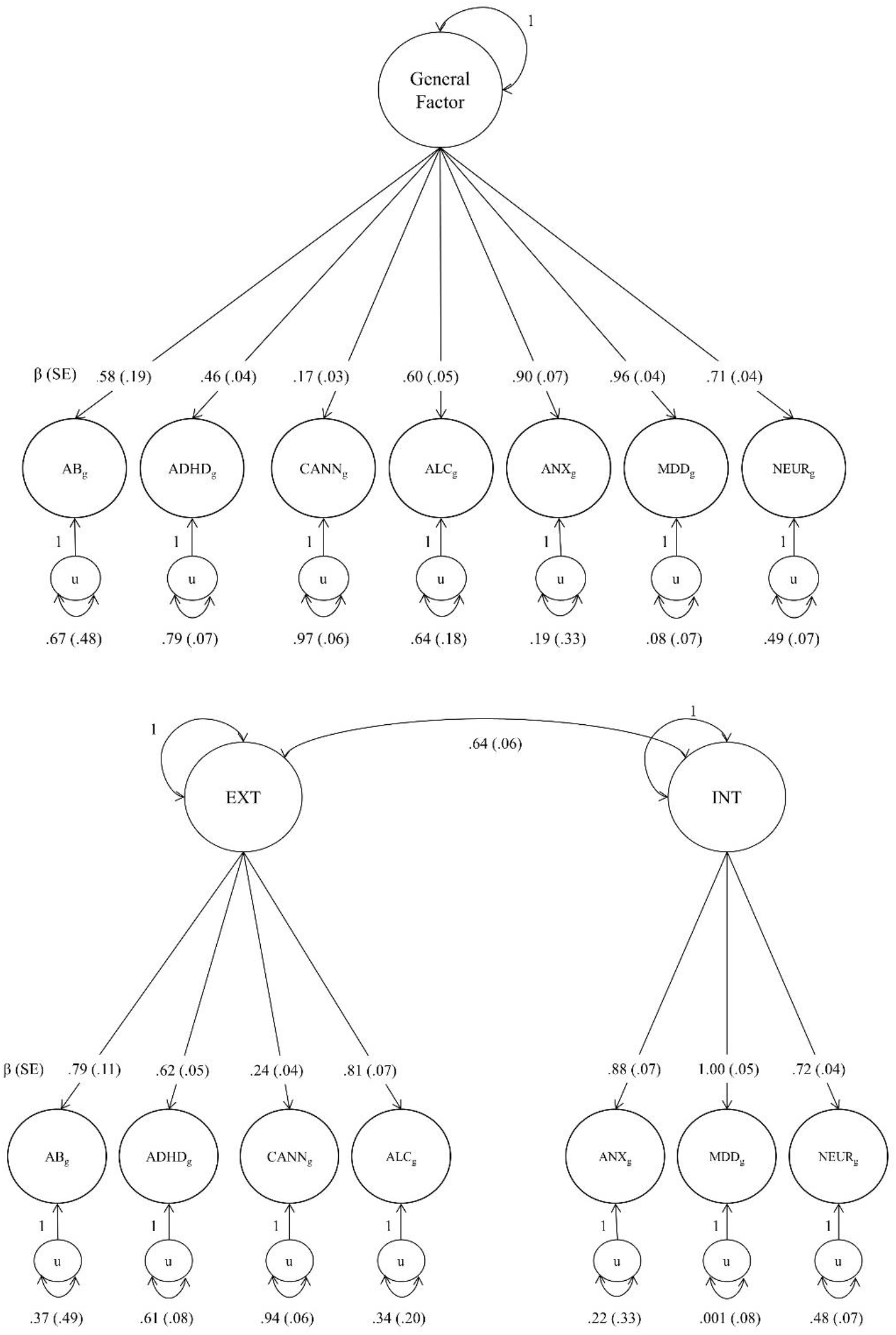
Genomic SEM one-factor and two-factor model. *Note*. Confirmatory factor analyses were conducted on the GWAS summary statistics in Table 1, using the Genomic SEM package in R Statistical Software (Grotzinger et al., 2019). Standardized estimates are shown. See Supplemental Figure 3 for unstandardized estimates. In both the one-factor and two-factor models, the residual variance of MDD was fixed to zero. Model fit comparisons between the one-factor model (χ^2^ [14] = 76.762, p < .001, AIC = 104.762, CFI = .962, SRMR = .127) and two-factor model (χ^2^ [13] = 46.072, p < .001, AIC = 76.072, CFI = .980, SRMR = .084) indicated superior model fit of the two-factor model (Δχ^2^ = 30.69(1), *p* < .001, ΔCFI > .01, lower AIC). Single-nucleotide polymorphism effects were then integrated into the model to derive new SNP weights for construction of latent polygenic scores (see Supplemental Methods).

Associations between the latent PGSs and phenotypic outcomes indicated that the latent internalizing PGS and latent one-factor PGS explained 1% more variance in the general factor of psychopathology than any of the individual PGSs that were used to construct these latent measures of polygenic risk (i.e., R^2^ = 2% versus R^2^ = 1%). There were no differences in the predictive power of the latent internalizing PGS and the latent one factor PGS, as indicated by non-overlapping confidence intervals of the standardized effects. Pooling the summary statistics of the externalizing GWAS (i.e., alcohol dependence, cannabis use, ADHD, antisocial behavior) similarly resulted in novel associations with internalizing phenotypes and the general factor of psychopathology, as compared to any of the individual externalizing PGSs. However, the model R^2^ were < 1% and there were no associations between the latent externalizing PGS and any of the externalizing outcomes.

## Discussion

We evaluated both the phenotypic and polygenic structure of psychopathology in a large population-based sample of older adults. In models that replicated using a split-half design, phenotypes were organized in a one-factor model of psychopathology rather than the two-factor internalizing-externalizing structure more common in younger samples (10,13). The general factor of psychopathology was further equivalent across gender and age groupings as indicated by invariant factor structure and loadings, suggesting that the structure of psychiatric phenotypes in the HRS is replicable across demographic groups. PGS analyses revealed that genetic risk scores derived from GWAS of externalizing psychopathology are not portable to older adults in the HRS: none of the externalizing PGSs were associated with externalizing or internalizing phenotypes. By contrast, the internalizing PGSs were predictive of internalizing phenotypes and the general factor of psychopathology in the current sample. Perhaps most importantly, using Genomic SEM (29), we found that the latent internalizing PGS and the latent one-factor PGS explained 1% more variance than any of the individual PGSs in models predicting internalizing phenotypes and the general factor of psychopathology. Collectively, these results make important contributions to our understanding of transdiagnostic risk for psychopathology – at phenotypic and genetic levels of analysis. For researchers and clinicians interested in the etiology and course of psychopathology in older adults, modeling general psychopathology is likely to improve predictive accuracy and may be important in developing interventions to reduce the burden of mental illness in the second half of the lifespan.

In contrast to research in children and adults (10,13), psychiatric phenotypes in older adults organized into one general factor of psychopathology rather than a two-factor internalizing-externalizing factor structure. Identification of the meta-structure of psychiatric phenotypes in older adults has both etiological and clinical implications. First, the largest loading on the general factor was negative affect. Negative affect or negative emotionality is thought to be a nonspecific vulnerability factor for multiple forms of psychopathology (64), is correlated with both internalizing and externalizing disorders (65), and is oftentimes the first factor extracted from individual differences in dispositional traits (64,66,67). That negative affect as a dispositional construct is robustly associated with multiple symptom domains (13) supports the Research Domain Criteria (RDoC) framework from the National Institute of Mental Health, in which the biological origins of intermediate phenotypes are linked to multiple categorical disorders (68). Our results further support the Hierarchical Taxonomy of Psychopathology (HiTOP) approach (69), which advocates for dimensional approaches that better characterize psychiatric comorbidity across symptom domains compared to traditional categorical nosologies. Clinically, interventions designed for one disorder have widespread effects on multiple disorders within the same domain (70). For example, pharmacological and psychosocial interventions designed to treat depression are also effective in treating some forms of anxiety (71), which has led to transdiagnostic interventions for emotional disorders broadly (72).

One major contribution of our results is the lack of specificity in PGS prediction of psychiatric phenotypes. It is surprising that a PGS designed to capture genome-wide genetic risk for a single disorder (e.g., MDD) was no better at predicting within domain (e.g., depressive symptoms) than cross-domain (e.g., state anger) phenotypes. One explanation for these results is that psychiatric GWAS rarely account for comorbidity (e.g., MDD cases without comorbid Substance Use Disorder). By ignoring psychiatric comorbidity, GWAS may be identifying genetic risk factors for multiple phenotypes or clinical severity instead of a single phenotype. Examples of psychiatric genetic studies that account for comorbidity include a study of bipolar disorder and schizophrenia (73) and a GWAS of comorbid depression and alcohol dependence (74). Precision phenotyping of homogenous subgroups (e.g., stratification by age of disorder onset) is also likely to improve to GWAS and resultant PGSs (75,76).

Using Genomic SEM, polygenic risk organized into a two-factor internalizing-externalizing structure, although the one-factor model also fit the data well. Importantly, these latent PGSs that aggregated genetic effects across multiple GWAS explained 1% more variation in the general factor of psychopathology. As MDD was the largest loading in both the one-factor and two-factor Genomic SEM models (Figure 4), it may not be surprising that there were no differences in the predictive power of the latent internalizing PGS and the latent one-factor PGS. Collectively, our results reiterate the power of aggregating genetic effects across multiple related phenotypes (27) and suggest that any researcher interested in capturing genome-wide genetic risk for psychopathology should implement methods to aggregate GWAS summary statistics of similar phenotypes rather than rely on PGSs of individual disorders.

In addition to practical implications, our results demonstrate that genetic risk for psychiatric phenotypes is transdiagnostic. Psychiatric GWAS repeatedly show that associated SNPs tend to cluster in genes underlying neurodevelopmental processes, signal transduction, and synaptic plasticity (39,41,77), all processes common to complex diseases. Moreover, biometric analyses in behavioral genetic/family designs demonstrate that a general genetic factor influences multiple psychiatric disorders (and their overlap) and explains more of the variation in psychiatric outcomes than the unique internalizing and externalizing genetic effects (19,78). More research is needed to understand whether psychiatric polygenic risk is pleiotropic and if so, what kind of pleiotropic processes are at play. For example, biological pleiotropy would suggest that a genetic risk variant for neuroticism (or another intermediate transdiagnostic phenotype) predicts multiple disorders (79,80). By contrast, mediated pleiotropy would suggest that a genetic risk variant predicts one phenotype (e.g., neuroticism), which subsequently predicts the onset of other phenotypes (e.g., alcohol use). Longitudinal phenotypic data and causal inferences techniques (81) are needed to evaluate these hypotheses.

A second explanation for low polygenic specificity in the current study is that PGSs are derived from GWAS of common genetic variation – most often SNPs with minor allele frequencies greater than 1% (82). An ‘omnigenic model of complex traits’ suggests that SNPs that contribute to the bulk of heritability in complex disorders are spread across the genome as common variants of small effect that contribute to cellular processes (e.g., protein binding, sequence-specific DNA binding) are common to many complex disorders. Disease-specific genetic risk variants, by contrast, are likely to be rare variants of large effect that are often not captured in GWAS of common genetic variation (22,23). Moreover, GWAS do not capture copy number variants (CNVs), which are also linked to psychiatric disorders and may function in a disease-specific manner (83). Thus, it may also be that PGSs derived from GWAS of common genetic variants are not appropriate for examinations of disorder-specific etiology.

Collectively, these results challenge the notion of specificity in the phenotypic and genetic presentation of psychopathology in older adults. The still impairing rates of internalizing and externalizing disorders during the second half of the lifespan necessitate discussion regarding the clinical utility of current diagnostic categories, particularly as we investigate psychiatric etiology using biological approaches such as genetics and neuroscience.

### Limitations

Although the current study is the first to evaluate the meta-structure of phenotypic and genetic risk for psychopathology in older adults using a large, population-based sample, several limitations are worth noting. First, the estimation of latent factors in confirmatory factor analysis is dependent upon the quality of the indicators. Based on previous recommendations (48,84), we only constructed PGSs based on large GWAS meta-analyses with independent replication samples. As a result, we did not include PGSs derived from smaller GWAS of relevant phenotypes, including several studies of externalizing disorders (85,86). Relatedly, the phenotypic measures available in the HRS were abbreviated scales, as is common in large population-based surveys. Thus, one alternative phenotypic model that we were unable to fit is a bifactor model of psychiatric outcomes (our models didn’t converge, likely due to the sparse measurement of symptoms), which posits that there are internalizing and externalizing factors as well as a higher-order bifactor that captures shared variance between the lower-order factors (10,13,65). Indeed, we observed a high correlation between the internalizing and externalizing factors in our sample, which is thought to indicate the presence of a higher-order bifactor (10,13,88). Moreover, externalizing phenotypes are not well represented in the HRS, likely because behaviors like aggression and rule-breaking are less among older adults. Antisocial behavior in childhood further places individuals at risk of early mortality or long-term incarceration (89), suggesting that individuals with the highest levels of externalizing behavior may not be represented in the HRS. Nevertheless, externalizing disorders such as ADHD and Substance Use Disorder are still common: between 3 – 4% of adults aged 55 – 85 meet criteria for ADHD (90) and 3.8% of adults over aged 55 meet criteria for a Substance Use Disorder (6). The non-significant associations between polygenic risk and externalizing behaviors in the HRS may be due to limited measures (e.g., impulsivity, trait anger, state anger, number of drinks per day) that do not adequately capture the complexity of externalizing behaviors in this age group.

Second, the GWAS summary statistics that we used to construct PGSs did not exclusively focus on older adults. Although maximizing statistical power through increasing sample size is a key consideration in GWAS, PGSs constructed from GWAS in younger samples may not generalize to older adults. This is particularly relevant considering the negative association we observed between impulsivity in the current sample and the PGS of antisocial behavior, constructed from a GWAS of adolescents and young-to-middle age adults (45). GWAS of psychiatric outcomes in pediatric cohorts (86,91) are beginning to show that genetic risk alleles may vary by developmental stage. As the availability of genomic data increases, future research should consider age-stratified GWAS.

Third, our analyses only focused on a subset of the population: older U.S. adults of European ancestry. Though the focus on older adults is a critical addition to research on the meta-structure of psychiatric disorders in adulthood, psychiatric genetics and human genetics studies overall are overwhelmingly Eurocentric (92) – a trend that reduces generalizability of all genetic work and is likely to exacerbate health disparities (93). We did not include participants of African ancestry in the current study because the available GWAS were conducted in European samples and, thus, would not be comparable for methodological rather than substantive reasons.

### Conclusion

Using multiple genome-wide PGSs for psychiatric outcomes, validated phenotypic measures, and novel analytic techniques in a relatively large, population-based sample of older adults, we showed that a single general factor of psychopathology best explained the phenotypic meta-structure of psychopathology in older adults. Moreover, although individual PGSs were non-specific in their associations with internalizing and externalizing outcomes, latent PGSs that aggregated genetic effects across several disorders explained 2% of the variation in phenotypes and explained more transdiagnostic variation than any individual PGS. These results inform a changing conceptualization of psychiatric diagnoses and their genetic etiology – from disorder-specific to transdiagnostic and dimensional.

## Acknowledgements

This research was funded by the National Institutes of Health T32-HD007109 and R25 AG053227 (AMG), R01-AG055406 and R01-AG055654a (EBW, JDF), L60-MD012145 (EBW), K99-AG05659 (LS), P30-AG012846 (LS, EBW), a NARSAD Young Investigator award from the Brain and Behavior Foundation (LWH), R01 MH121079 (LWH, CM), and R01 MH110872 and R01 MD011716 (CM).

The Health and Retirement Study (HRS) is supported by the National Institute on Aging (U01 AG009740). HRS genotyping was funded separately by the National Institute on Aging (RC2 AG036495, RC4 AG039029) and was conducted by the NIH Center for Inherited Disease Research (CIDR) at Johns Hopkins University. Genotyping quality control and final preparation of the genotype data were performed by the University of Michigan School of Public Health and the Genetics Coordinating Center at the University of Washington.

## Supplemental Methods

### Genetic Ancestry Estimation

SNPs used for PCA were selected by LD pruning from an initial pool consisting of all autosomal SNPs with a missing call rate < 5% and minor allele frequency (MAF) > 5%, and excluding any SNPs with a discordance between HapMap controls genotyped along with the study samples and those in the external HapMap data set. In addition, the 2q21 (LCT), HLA, 8p23, and 17q21.31 regions were excluded from the initial pool (1).

Additional information about the genetic quality-control procedures in the Health and Retirement Study can be found at https://hrsonline.isr.umich.edu/sitedocs/genetics/HRS2_qc_report_SEPT2013.pdf?_ga=2.59963114.6513449.1600955376-325044460.1531409717&_ga=2.59963114.6513449.1600955376-325044460.1531409717

## Supplemental Results

### Measurement Invariance by Gender

Women reported greater negative affect (t[5923] = 7.31, *p* < .001), anxiety (t[5928] = 3.77, *p* < .001) and depressive symptoms (t[5965] = 7.02, *p* < .001), and men reported greater state anger (t[5927] = 5.05, *p* < .001). Despite mean level gender differences, the one-factor model of general psychopathology fit well in both men and women (Figure 3a). Measurement invariance testing revealed metric invariance across genders: fixing the indicator loadings to be equivalent across groups did not significantly degrade model fit. Not surprisingly, given mean differences in the individual measures by gender, the model did not meet criteria for scalar measurement invariance (i.e., equivalent intercepts across groups), indicated by ΔCFI = .031 and ΔTLI = .015.

### Measurement Invariance by Age

Middle age (51 – 64 years; *n* = 2,212), young-old (65 – 74, *n* = 2,366), and old-old (75 – 83, *n* = 1,425) HRS participants significantly differed in their mean levels of negative affect (F[2,5922] = 47.64, *p* < .001), trait anger (F[2,5935] = 44.72, *p* < .001), state anger (F[2,5926] = 22.003, *p* < .001), impulsivity (F[2,5918] = 3.04, *p* < .05), and depressive symptoms (F[2,5998] = 73.33, *p* < .001). Tukey post-hoc comparisons revealed that middle age participants reported greater negative affect, trait anger, state anger, and depressive symptoms than young-old participants (all *p*s < .01). Young-old participants, in turn, reported greater trait and state anger than old-old participants, but did not differ from old-old participants on negative affect or depressive symptoms. Lastly, although middle-age and young-old participants did not differ on self-reported impulsivity, old-old participants reported greater impulsivity than both younger age groups. Despite mean-level differences in phenotypic outcomes by age group, the one-factor model of general psychopathology fit well in all three age groups (Figure 3b). As in invariance testing by gender, metric (i.e., fixed loadings), but not scalar (i.e., fixed intercepts), invariance was established across age groups (Figure 3b).

**Supplemental Table 1.** Descriptive statistics and sources of phenotypic measures

| Phenotypic Measures of Psychopathology |  |  |  |  |  |
| --- | --- | --- | --- | --- | --- |
| Construct | Mean (SD) | Min – Max | Reliability ($\alpha$) | Number of Items | Source |
| Negative Affect | 1.74 (.60) | 1 – 4.83 | .97 | 12 | Positive and Negative Affect Schedule – Expanded Form (2) |
| Anxiety Symptoms | 1.51 (.55) | 1 – 4 | .97 | 5 | Beck Anxiety Inventory (3) |
| Depressive Symptoms | 1.12 (1.76) | 0 – 8 | .98 | 8 | Center for Epidemiologic Studies Depression Scale (4) |
| Impulsivity | 2.69 (.83) | 1 – 6 | .97 | 6 | Multidimensional Personality Questionnaire (5) |
| Trait Anger | 2.21 (.69) | 1 – 4 | .96 | 4 | State-Trait Anger Expression Inventory (6) |
| State Anger | 1.49 (.49) | 1 – 4 | .97 | 7 | State-Trait Anger Expression Inventory (6) |
| Drinking Frequency* | .83 (1.41) | 0 – 24 | - | 1 | - |

Note. $5,921 > N > 6,001$ . All constructs except drinking frequency were measured as means of all items.
\*Participants were asked whether they currently consume alcoholic beverages and, if so, how many per day on average. A continuous measure was constructed, where non-drinkers were recoded as “0”.

**Supplemental Figure 1.**
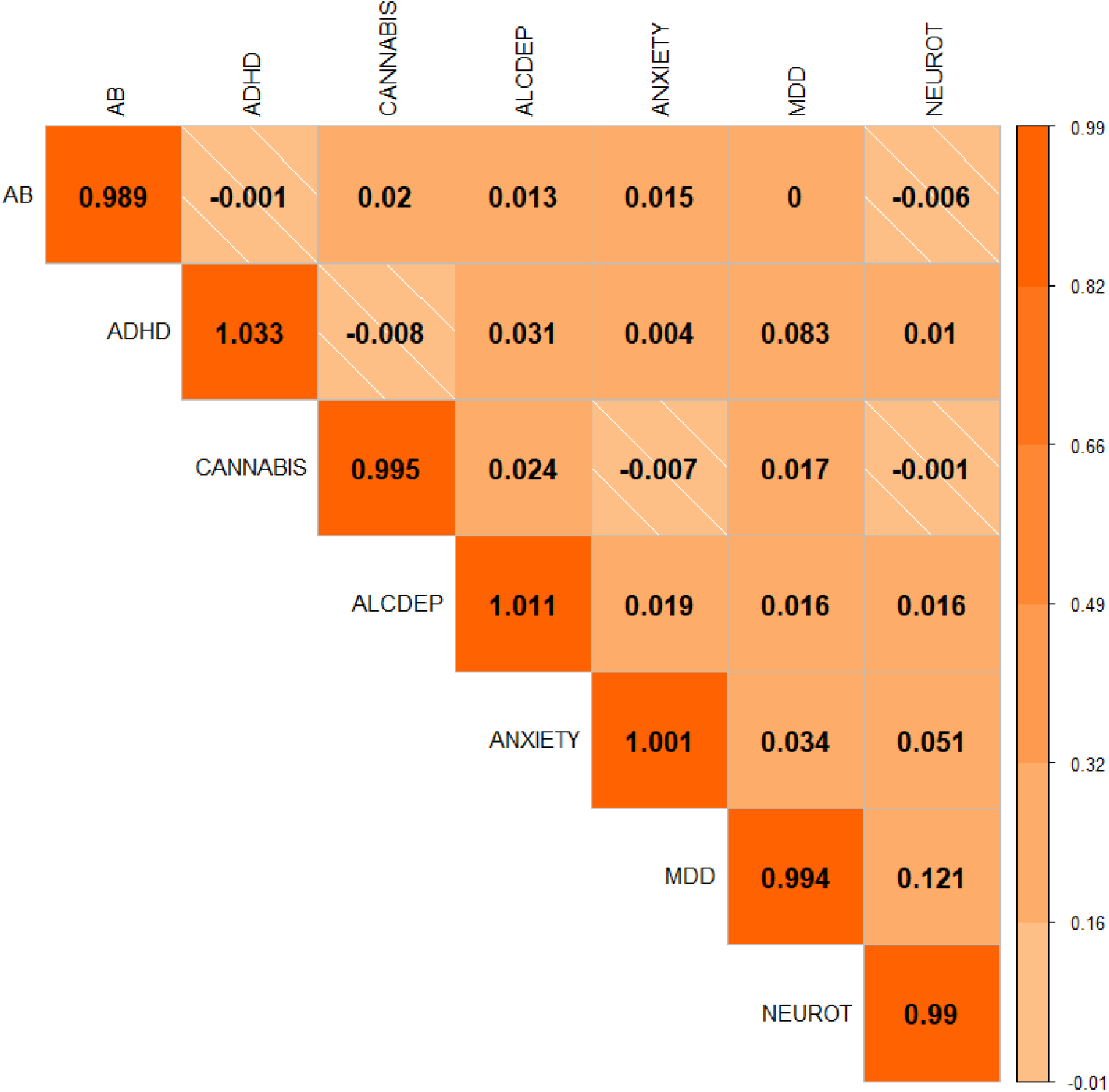
Non-zero cross-trait LD-score regression intercepts suggest cryptic relatedness among GWAS samples.

**Supplemental Figure 2.**
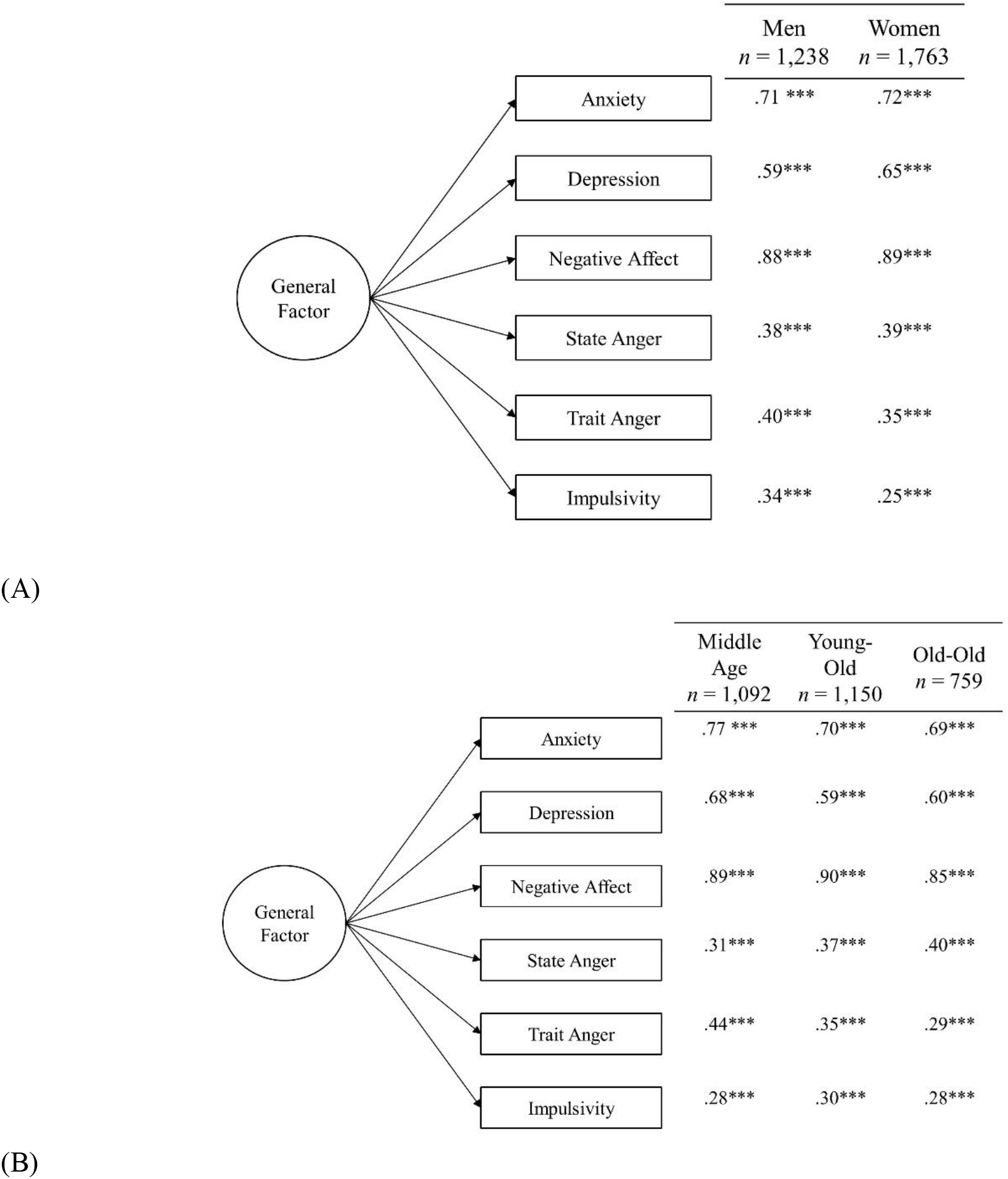
Invariant indicator loadings of the general factor of psychopathology by age group and gender in the Health and Retirement Study. Note. (A) Confirmatory one-factor phenotypic models in the hold-out sample, split by gender (model fit: χ^2^ (18) = 64.22, *p* < .001; CFI = .985; TLI = .975; RMSEA = .045, 90% CI [.033, .057]). Increasingly stringent measurement invariance testing revealed no significant change in model fit when loadings were fixed across groups (metric invariant model ΔCFI = .009, ΔRMSEA = 004), but a significant depreciation of model fit when loadings and intercepts were fixed across groups (scalar invariant model ΔCFI = .018, ΔRMSEA = .009). (B) Confirmatory one-factor phenotypic models in the hold-out sample, split by age group (model fit: χ^2^ (27) = 82.77, *p* < .001; CFI = .983; TLI = .971; RMSEA = .049, 90% CI [.037, .061]). Increasingly stringent measurement invariance testing revealed no significant change in model fit when loadings were fixed across groups (metric invariant model ΔCFI = .009, ΔRMSEA = .003), but a significant depreciation of model fit when loadings and intercepts were fixed across groups (scalar invariant model ΔCFI = .031, ΔRMSEA = .015).

**Supplemental Figure 3.**
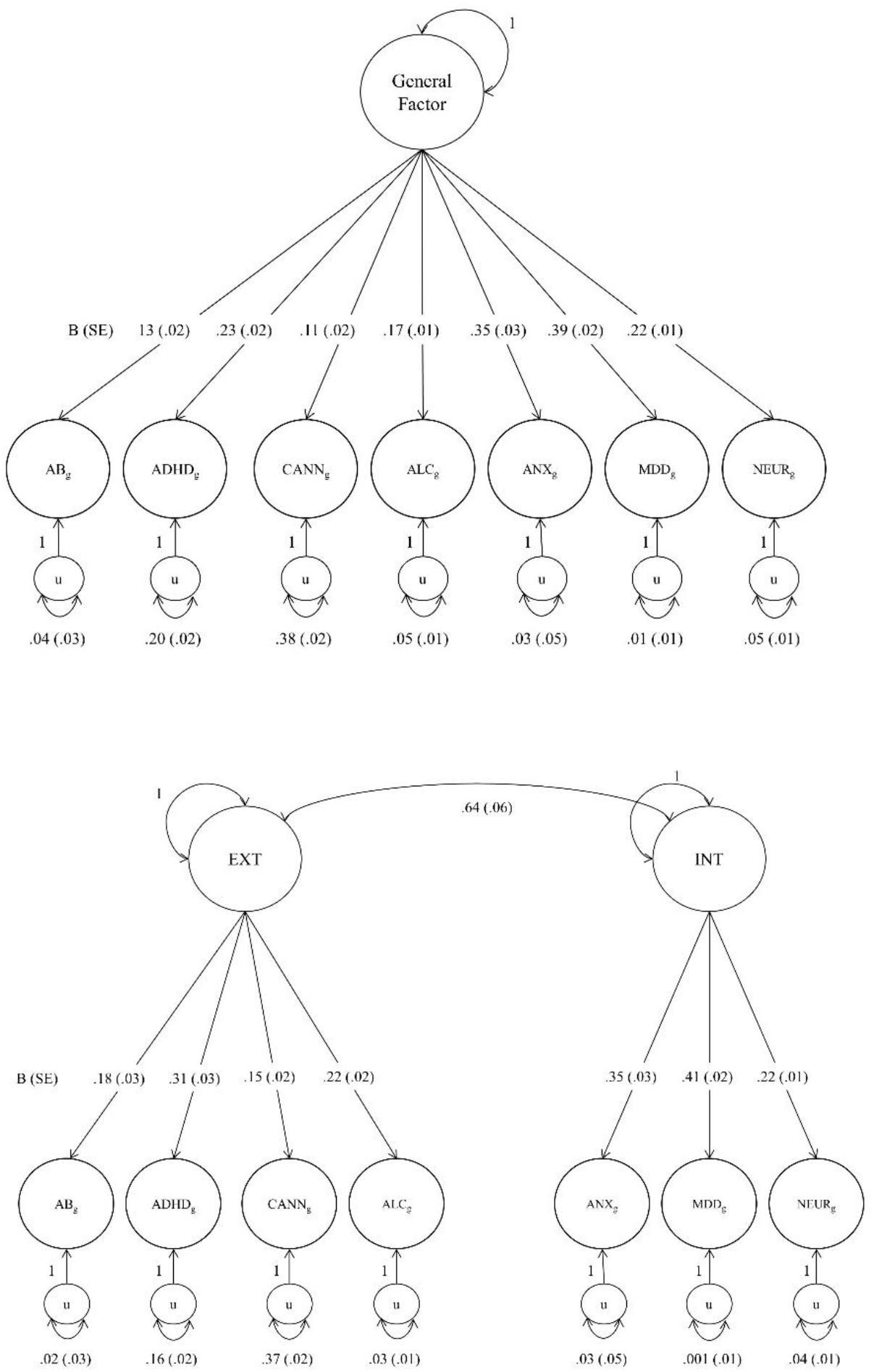
Unstandardized factor loadings for Genomic SEM one-factor and two-factor models. *Note*. Confirmatory factor analyses were conducted on the GWAS summary statistics in Table 1, using the Genomic SEM package in R Statistical Software (7). In both the one-factor and two-factor models, the residual variance of MDD was fixed to zero. Model fit comparisons between the one-factor model (χ^2^ [14] = 76.762, p < .001, AIC = 104.762, CFI = .962, SRMR = .127) and two-factor model (χ^2^ [13] = 46.072, p < .001, AIC = 76.072, CFI = .980, SRMR = .084) indicated superior model fit of the two-factor model (Δχ^2^ = 30.69(1), p < .001, ΔCFI > .01, lower AIC). Single-nucleotide polymorphism effects were then integrated into the model to derive new SNP weights for construction of latent polygenic scores (see Supplemental Methods).

